# Uttroside B, a US-FDA-Designated Orphan Drug Against Hepatocellular Carcinoma (HCC), Impedes Non-alcoholic Steatohepatitis (NASH) and NASH -Induced HCC

**DOI:** 10.1101/2024.11.06.622394

**Authors:** Tennyson P Rayginia, Chenicheri K. Keerthana, Sreekumar U. Aiswarya, Sadiq C. Shifana, S Jannet, Sanjay Suresh Varma, Joy P Maria, Mundanattu Swetha, Archana Payickattu Retnakumary, Lekshmi R. Nath, Kalishwaralal Kalimuthu, Vishnu Sunil Jaikumar, Sankar Sundaram, Nikhil Ponnoor Anto, Noah Isakov, Kuzhuvelil B. Harikumar, Ravi S. Lankalapalli, Ruby John Anto

**Affiliations:** Division of Cancer Research, Rajiv Gandhi Centre for Biotechnology, Thiruvananthapuram 695014, Kerala, India; Department of Biotechnology, University of Kerala, Thiruvananthapuram 695011, Kerala, India; Molecular Bioassay Laboratory, Institute of Advanced Virology, Bio360 Life Science Park, Thiruvananthapuram, 695317, Kerala, India; Chemical Sciences and Technology Division, CSIR-National Institute for Interdisciplinary Science and Technology, Thiruvananthapuram 695019, Kerala, India; Department of Pharmacognosy, Amrita School of Pharmacy, Amrita Vishwa Vidyapeetham, AIMS Health Science Campus, Kochi 682041, Kerala, India; Department of Pathology, Government Medical College, Kottayam 686008, Kerala, India; Centre de Recherche l’Hôpital Maisonneuve-Rosemont (CR-HMR), and Département de microbiologie, infectiologie et immunologie, Faculté de médecine, Universite de Montreal, QC, Canada; The Shraga Segal Department of Microbiology, Immunology and Genetics, Faculty of Health Sciences, Ben-Gurion University of the Negev, P.O. Box 653, Beer Sheva 84105, Israel; Centre of Excellence in Nutraceuticals, Bio360 Lice Science Park, Thiruvananthapuram, 695317, Kerala, India

**Keywords:** Uttroside B, NASH, HCC, NAFLD, Lipogenesis, Fibrogenesis

## Abstract

**Introduction:** Non-alcoholic steatohepatitis (NASH) is characterized by excessive accumulation of fat, accompanied by inflammation and liver injury. NASH can lead to chronic conditions like fibrosis and cirrhosis, and has an elevated risk of progressing to hepatocellular carcinoma (HCC). Currently there are no FDA-approved drugs for the treatment of NASH.

**Objectives:** Our discovery of Uttroside B (Utt-B), a phytosaponin isolated from *Solanum nigrum* Linn., which exhibits remarkable anti-HCC potential, has gained global recognition and is currently a US-FDA-designated ‘orphan drug’ against HCC. The present study highlights Utt-B as an anti-NASH molecule, by utilizing a High-Fat-Diet murine model, and as an inhibitor to the progression of NASH to HCC, using a streptozotocin-induced steatohepatitis-derived HCC animal model, thereby warranting its further validation as a propitious candidate drug molecule against NASH and NASH-induced HCC.

**Methods:** High fat diet-induced NASH and streptozotocin-induced steatohepatitis-derived HCC were developed in C57BL/6 mice. Utt-B was administered intraperitoneally. q-PCR, immunoblotting and staining techniques such as Haematoxylin and eosin, Oil Red O, Sirius Red and Masson’s Trichrome, were performed to assess the therapeutic potency of Utt-B against NASH. Nanostring n-Counter analysis was conducted to verify the anti-fibrotic potential of Utt-B in NASH-induced HCC mouse model.

**Results:** Utt-B ameliorates the pathological features such as, steatosis, hepatocyte ballooning and inflammation associated with NASH. Utt-B up-regulates the expression of autophagy markers ATG7, Beclin-1 and LC-III and down-regulates the expression of α-SMA, the indicator protein for the activation of hepatic stellate cells. Utt-B hinders the development of fibrosis and halts the progression of NASH to HCC in NASH-induced HCC mouse model.

**Conclusion:** Our investigation reveals that Utt-B effectively alleviates NASH and abrogates its progression to HCC. As no treatment options are currently available against NASH, our findings are very relevant and strengthen the prospect of developing Utt-B as a potent drug for the treatment of NASH and NASH-induced HCC.

**Graphical Abstract:** 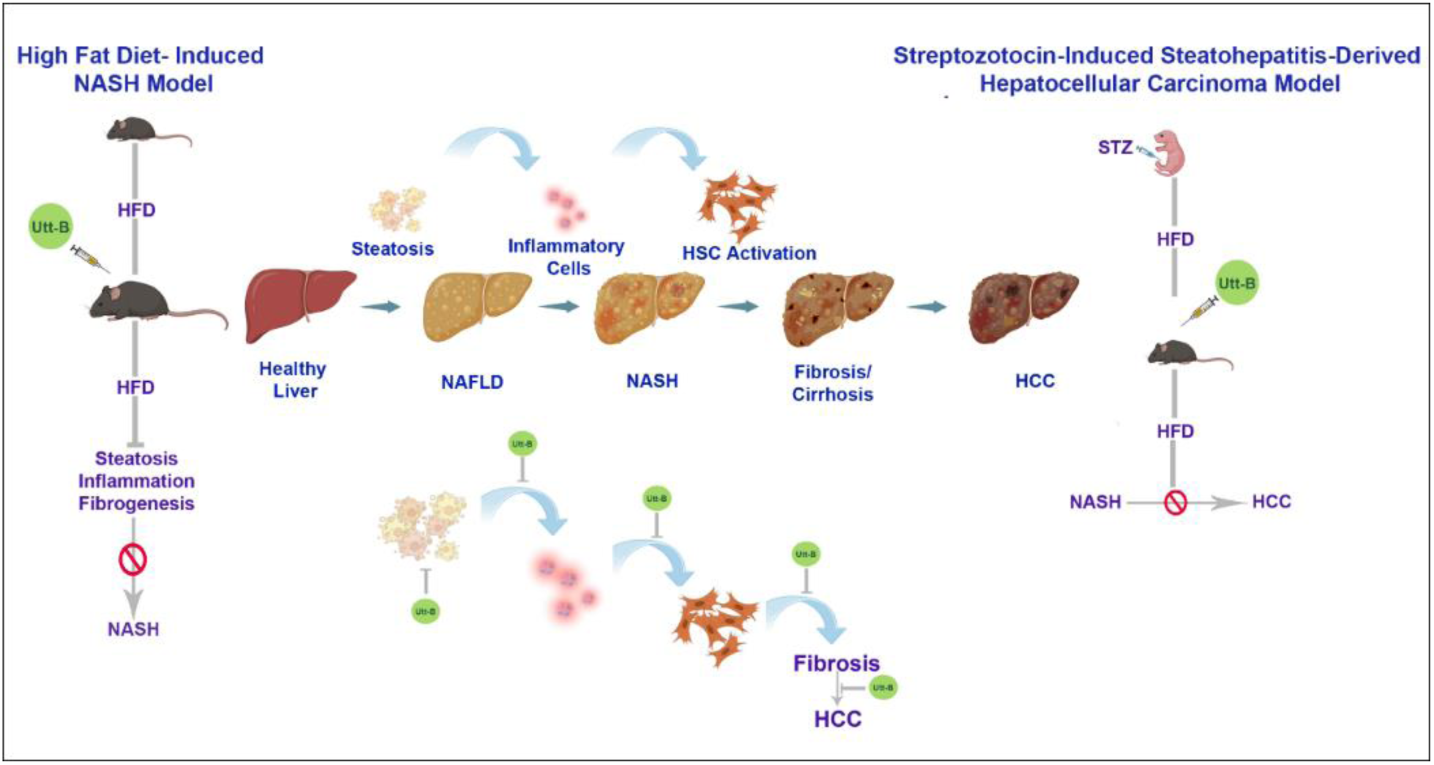

## Introduction

Non-alcoholic steatohepatitis (NASH) ensues simple hepatic steatosis in non-alcoholic fatty liver disease (NAFLD), a condition impacting approximately 40% of the adult population. The prevalence of NAFLD and NASH, has shown a notable increase in recent decades. While non-alcoholic fatty liver (NAFL) represents a relatively benign condition, NASH emerges as a more aggressive form marked by steatosis, inflammation, hepatocyte ballooning, hepatocellular injury, and fibrosis. Various detrimental factors, including lipotoxicity, pro-inflammatory signals and endotoxins, contribute to the pathophysiology of NASH. Further, prolonged liver injury in NASH significantly elevates the risk of advanced liver diseases, including cirrhosis and liver cancer [1–5].

Liver fibrosis is a major risk factor for the development of liver cancer in individuals with NAFLD [6]. Cytokines and chemokines released due to lipotoxic damage in hepatocytes, activate innate and adaptive immune cells, leading to an inflammatory cascade [7]. Damaged hepatocytes release extracellular vesicles comprising of apoptotic bodies, exosomes, and microparticles, that initiate the activation of non-parenchymal cells and the recruitment of immune cells, ultimately resulting in the progression of fibrosis [8, 9]. Hepatic injury resulting from lipogenesis and inflammation induces the activation of hepatic stellate cells (HSCs), thereby increasing the expression of α–SMA, TGFβ, and Collagen type 1, and promoting epithelial-to-mesenchymal transitions, which collectively contribute to the development of fibrosis [10–12].

Recent therapeutic advancements in the treatment of NASH encompass strategies targeting PPARs, FXRs, THRs, GLP-1, FGFs, and more. Innovations in the field also include stem cell-based therapy, interventions targeting ferroptosis, utilization of apoptosis inhibitors, microRNA treatment, strategies addressing oxidative stress, and modulation of the gut microbiome for NASH therapy. Despite ongoing research efforts, the precise pathogenic mechanisms of NASH remain elusive and hence there is a lack of effective pharmaceutic interventions against NASH, except the recent US-FDA-approved small molecule drug, resmetirom. Although there has been notable progress in developing disease management strategies focused on attenuating fibrosis, NASH is poised to emerge as a prominent contributor to the development of hepatocellular carcinoma (HCC) [5].

Previous investigations by our research team, have highlighted the remarkable efficacy of Uttroside B (Utt-B), a phytosaponin isolated in our lab, from the methanolic extract of the leaves of *Solanum nigrum* Linn. *(S. nigrum)*, against HCC. This groundbreaking innovation was granted patent by the USA (US20190160088), Canada (3,026,426), Japan (JP2019520425), South Korea (KR1020190008323) and Europe (EP3463382). Additionally, Utt-B also received the ‘Orphan Drug’ designation against hepatocellular carcinoma from the US FDA [13–15]. Several studies have reported the anti-tumor, anti-inflammatory, anti-oxidant, hepatoprotective and immunomodulatory efficacy of *S. nigrum* [16–20]. Given the exceptional efficacy of Utt-B, an active component of *S. nigrum*, against hepatocellular carcinoma (HCC), the present study has explored the therapeutic potential of Utt-B in the context of NASH and NASH-induced HCC by employing two different murine models; High Fat Diet-induced (HFD) NASH model and Streptozotocin-induced steatohepatitis-derived hepatocellular carcinoma model (STAM). Both models replicate the pathological features associated with human NASH and NASH-induced HCC, thus providing comprehensive pre-clinical data that demonstrate Utt-B as a potent molecule, which possesses all the credentials to be evaluated as a candidate drug for the treatment of NASH as well as NASH-induced HCC.

## Materials and Methods

### Materials

A High Fat Diet containing 60% kcal fat was purchased from National Institute of Nutrition (NIN), Hyderabad, India. Reagents used for staining mouse liver tissues such as Haematoxylin (H9627), Eosin (230251), Oil Red O (O0625), and Direct Red (Sirius Red) (365548) were obtained from Sigma Chemicals (St.Louis, MO, USA). Trichrome staining kit (ab150686) was purchased from Abcam. Antibodies against, ATG-7 (2613P), Beclin-1 (3495P), LC-III (4599P), ACC (3662S), GAPDH (2118S), β-actin (12620S) and Vinculin (13901S) were obtained from Cell Signalling Technologies (Beverly, MA, USA) and the antibodies against α-SMA (sc-53142) and phospho-ACCα (sc-271965) were purchased from Santa Cruz Biotechnology (Santa Cruz, CA, USA). cDNA synthesis kit and SyBr Green for qRT-PCR was procured from Applied Biosystems (4367659). Poly Excel HRP/DAB detection system universal kit (PathnSitu Biotechnologies Pvt. Ltd., Secunderabad, India, OSH001) was used for immunohistochemistry experiments. All other chemicals were purchased from Sigma Chemicals (St.Louis, MO, USA) unless otherwise mentioned.

## Methods

### Isolation and purification of Utt-B

Isolation and purification of Utt-B were done as previously reported [13].

### Ethics Statement

All experiments using laboratory animals were carried out in compliance with relevant laws and institutional guidelines of RGCB. The experiments were conducted after obtaining prior approval from the Institutional animal ethical committee of RGCB (IAEC) (IAEC/899/2022 and IAEC/853/RUBY/2021) and have followed the rules and regulations prescribed by the Committee for Control and Supervision of Experiments on Animals (CCSEA), Government of India.

### In vivo Experiments

#### High-Fat-Diet Mouse Model

High fat diet induced NASH was developed in C57BL/6 mice as previously described with slight modifications [48, 49]. Seven weeks old, male C57BL/6 mice (3-5 per cage), were housed in a 12-hour light/dark cycle, for a total duration of six months. Following one week of acclimatization with a standard chow diet and water, the animals were randomly divided into three experimental groups (n=6, per group), namely Control, HFD and HFD+Utt-B groups. Control group received the standard chow diet; HFD mice were fed HFD for 6 months; Mice in the HFD+Utt-B group were fed a HFD and treated with Utt-B at a dose of 10mg/kg, on alternate days, starting from the 3^rd^ month of consumption of HFD to the 6^th^ month. The mice fed a HFD (HFD and HFD+Utt-B groups) were injected intraperitoneally with 5 doses of T0901317 and 4 doses of CCl_4_ during the 6^th^ month, as described previously [48, 49]. Corn oil served as a vehicle for the CCl_4_ and T090137 (T2320, Sigma) injections. After the duration of the study, the animals were euthanized following which, blood and tissue samples were collected for further analyses.

#### Streptozotocin-Induced Steatohepatitis Derived Hepatocellular Carcinoma (STAM) Mouse Model

STAM mice were developed as previously described [21–23]. 2 days old C57/BL6 pups were injected with 200µg of streptozotocin in sodium citrate buffer. After weaning the mice were separated according to gender, and both male and female mice were used in the study. At 4 weeks of age, the control mice, which were injected with sodium citrate buffer alone at 2 days of age, were fed a standard chow diet, and the mice in STZ+HFD and treatment groups were fed HFD (n=6, per group). STAM mice have been reported to develop NASH by 8th week, fibrosis by 12-13^th^ week and HCC by 20^th^ week. Hence, animals in the treatment group were administered 10mg/kg of Utt-B in PBS starting from the 8^th^ week, the stage at which NASH has been fully developed. The duration of study in the male mmice was for a period of 22 weeks whereas the female mice were caged for approximately 30 weeks, as it was reported that the female hormones would interefere with the action of streptozotocin. Accordingly, Utt-B treatment was continued until the 22^nd^ week in the male mice whereas drug treatment was continued until the 30^th^ week in the female mice. After the duration of the study, the animals were euthanized following which, blood and tissue samples were collected for further analyses.

#### Biochemical Analysis

Serum biochemical assays were conducted for the analysis of serum total cholesterol, triglycerides, creatinine, AST and ALT was performed in Fujifilm Full Dri Chem NX500i.

#### Oil Red O (ORO) staining

The stock solution of ORO was prepared by dissolving 25g of ORO in 400ml of 99% Isopropyl alcohol and mixing by keeping on a magnetic stirrer for 2hrs at room temperature. To prepare working stock, ORO stock and distilled water were mixed in a ratio of 1.5:1 and filtered through a 0.45μm filter. Oil Red O working solution was added dropwise to tissue section obtained from frozen liver tissue and was incubated for 5 minutes. It was counterstained with haematoxylin for 15 seconds, followed by rinsing in tap water for 30 minutes. After checking whether staining is sufficient or not under the microscope, the sections were mounted using glycerol and sealed along the edges with nail polish [50]. Images were captured utilizing a DMi8 Inverted Fluorescence Research Microscope, complemented by a DMC 2900 Digital Camera.

#### Sirius Red Staining

Formalin-fixed paraffin-embedded tissue sections were subjected to a standardized deparaffinization protocol. Initially, they were immersed in Xylene to eliminate paraffin residues. Subsequently, the sections were sequentially treated with 100%, 95%, and 70% Isopropyl alcohol (IPA) for precise durations of 8, 5, and 5 minutes, respectively. Sirius Red staining was done according to the manufacturer’s protocol. Following this, a thorough rinse with distilled water was conducted for 5 minutes, followed by staining with Mayer’s Hematoxylin for a defined period of 30 minutes. Excess stain was carefully removed through immersion in distilled water, succeeded by a brief exposure to acid alcohol. Subsequent steps involved a tap water rinse prior to bluing, which entailed a precisely timed incubation of 10 minutes in bluing solution. Application of Sirius red (Direct Red) stain ensued, with meticulous attention paid to the dropwise addition onto tissue sections, followed by an incubation period of 1 hour. Acidified water washes were meticulously performed twice, each at equal intervals of 5 minutes. Upon thorough air-drying, slides underwent two consecutive washes in 100% ethanol, followed by a final immersion in xylene. Ultimately, the sections were meticulously mounted in DPX and subjected to examination under a light microscope. Images were captured utilizing a DMi8 Inverted Fluorescence Research Microscope, complemented by a DMC 2900 Digital Camera.

#### Hematoxylin and Eosin (H&E) Staining

Tumor and liver tissue samples derived from mice were subjected to fixation, sectioning, and subsequent staining employing Hematoxylin and eosin staining [14]. The samples were blinded before being analysed by the pathologist. Images were captured utilizing a DMi8 Inverted Fluorescence Research Microscope, complemented by a DMC 2900 Digital Camera.

#### Trichrome Staining

The staining was performed according to the manufacturer’s protocol using the Trichrome staining kit (Abcam), a kit designed to visualize connective tissues, more particularly collagen in tissue sections, strictly adhering to manufacturer’s protocol. The tumor and liver tissue sections were deparaffinized by being dipped in xylene. Then, Bouin’s Fluid was preheated to a temperature range of 54-64°C. Next, the slide was incubated in the preheated Bouin’s Fluid for 60 minutes, followed by a cooling period of 10 minutes. The slide was rinsed in water before being incubated in Weigert’s Iron Hematoxylin for 5 minutes. After another water rinse, the slide was immersed in Biebrich Scarlet/Acid Fuchsin solution for 15 minutes. This was followed by differentiation in a phosphomolybdic/phosphotungstic acid solution for 10-15 minutes. Subsequently, the slide was incubated in Aniline Blue solution for 5-10 minutes before being rinsed again in water. Then, the slide was immersed in an acetic acid solution for 3-5 minutes. Finally, the process was completed by dehydrating, clearing, and mounting the slide. Images were captured utilizing a DMi8 Inverted Fluorescence Research Microscope, complemented by a DMC 2900 Digital Camera.

#### Immunohistochemistry

Immunolocalization of specific proteins within tissue sections was conducted utilizing the Poly Excel HRP/DAB detection system, a universal kit designed for mouse and rabbit primary antibodies (OSH001, PathnSitu Biotechnologies Pvt. Ltd, India), adhering strictly to the manufacturer’s protocol. All immunohistochemistry images were captured utilizing a DMi8 Inverted Fluorescence Research Microscope, complemented by a DMC 2900 Digital Camera.

#### Immunoblotting

Lysates were prepared in whole lysis buffer, followed by centrifugation for 20 minutes at 14,000 rpm at 4°C, and the supernatant was collected. Protein concentration was estimated using Bradford’s method. Subsequently, 30 μg of protein was loaded per well, and proteins were separated through SDS-polyacrylamide gel electrophoresis (PAGE) before being electrophoretically transferred to polyvinyl difluoride (PVDF) membranes in 20% methanol TRIS-glycine transfer buffer. After blocking with 5% skimmed milk in TBST, membranes were probed using antibodies as specified above.

#### RNA isolation

RNA isolation from mouse liver tissues was performed by TriZol method. Liver tissue weighing 10-15µg was lysed with 1000µl trizol, passed through a syringe and vortexed once. The tissue lysate was incubated for 5 minutes at room temperature. 0.2 millilitre of chloroform was added and vortexed vigorously for 15 seconds and then kept for incubation at room temperature for 2-3 minutes. Further the samples were centrifuged at 12000*g for 15 minutes at 2-8°C. The aqueous phase was removed and 0.5ml of isopropanol was added, mixed gently and incubated at 15-30°C for 10 minutes. Further centrifugation was carried out at 12000*g for 10 minutes at room temperature. The RNA precipitate often invisible before centrifugation formed a gel like precipitate on the side and bottom of the tube. The entire supernatant was gently removed. The RNA pellet was washed with 75% ethanol. The samples were further mixed by vortexing and centrifuged at 7500*g for 5 minutes at 2-8 °C. The washing step was repeated to remove all the left-over ethanol. Finally, the RNA pellet was air dried for about an hour. The purity and yield (in ng/μl) of the mRNA samples were measured using Nanodrop. The RNA samples were stored at −80°C until and after use.

#### cDNA synthesis

cDNA was synthesized from the respective RNA samples using Applied Biosystems cDNA synthesis kit.

#### Real Time RT-PCR

The stored cDNA was thawed and diluted (1:10) with nuclease free water. The SyBr Green reagent from Applied Biosystem was used for the experiment. Ten microlitre of the reaction mix (2.5 μl of cDNA +7.5 μl of master mix) was added in triplicates for genes MCP-1, IFN-γ, TNF-α, RCP, Collagen1α and PPARγ. GAPDH was added to the wells of a 96 well PCR plate and placed in a thermal cycler. The ddCT fold change in the gene levels were analysed. The statistical significance of the PCR results was analysed using Graph Pad Prism software version 8.0.

#### NanoString mRNA Expression assay

For NanoString analysis, samples were quantified using Qubit RNA HS assay (Invitrogen, Cat # Q32855) kit and also qualitatively analysed on Agilent 2100 bioanalyzer nano chip (Agilent, Cat # 5067-1511). Codeset reaction for nCounter Mouse Fibrosis v2 Panel (NanoString Technologies) was carried out as per the Manual (nCounter Gene Expression Panel and custom codeset user manual, MAN-10056-06). RNA was hybridized with Reporter and Capture probes at 65degC for 18 hours and samples run on nanoString nCounter SPRINT machine (1) through service provider Theracues Innovations Pvt Ltd, Bangalore, India. The Raw data files were downloaded from nCounter SPRINT machine and analyzed using nSolver™ 4.0 (NanoString Technologies) software.

The foldchange values and following un-paired Student’s T-test based pvalue were calculated using the ‘calculate ratio’ module from nSolverTM 4.0 Analysis Software (MAN-C0019-08). Differentially expressed genes were identified with thresholds of pvalue < 0.05 and Foldchange > 1.2 (upregulated) or Foldchange < −1.2 (downregulated). The pvalue was corrected using the FDR approach. The genes were identified if expressed above background threshold on the basis of their normalized expression value if expressed above the average of negative probe counts. All statistical analysis was performed by using R (version 4.0.2, https://www.r-project.org/). P value below 0.05 was considered statistically significant.

#### Statistical analysis

Statistical analyses were performed using GraphPad Prism software Version 8 (GraphPad Software Inc., San Diego, CA, USA). To assess statistical significance, Student’s t-tests, One-way ANOVAs, and Two-way ANOVAs were employed, with a significance threshold set at p < 0.05. The error bars represent ± SD and are indicative of three independent experiments.

## Results

### Utt-B inhibits development of high-fat diet (HFD)-induced NASH

As Utt-B exhibited remarkable anti-HCC potential in our previous studies [13–15], we investigated the efficacy of the compound against NASH, another major liver disease, the progress of which may lead to the onset of HCC. NASH was induced in C57BL6 mice by administering HFD for a duration of 6 months, supplemented with intraperitoneal injections of CCl_4_ and T0901317, as described in the Materials and Methods. It was observed that NASH-related features such as steatosis and ballooning of hepatocytes start to develop in HFD-mice after 3 months of consumption of HFD (Fig S1A). Hence, to assess the potential therapeutic impact of Utt-B against NASH, animals in the treatment group were administered with 10mg/kg of Utt-B intraperitoneally in PBS, from the 4^th^ month onwards (Fig 1A). A steady rise in the body-weight of animals in the HFD group was noted compared to those fed with a standard chow diet, while the treatment group exhibited a progressive reduction in body-weight, post-Utt-B administration. Morphologically, the livers resected from mice in the HFD group were enlarged and pale, with a rough texture, in contrast to the normal-sized livers with a smooth texture, as observed in both the control and Utt-B treated groups (Fig 1B). This morphological variation was corroborated by the liver-to-body weight ratio, which differed among the three groups. The liver-to-body weight ratio in the HFD group was markedly higher than that in the control, whereas that of Utt-B treated mice exhibited a significant reduction and was comparable with that of the contro (Fig 1C)

**Fig 1:**
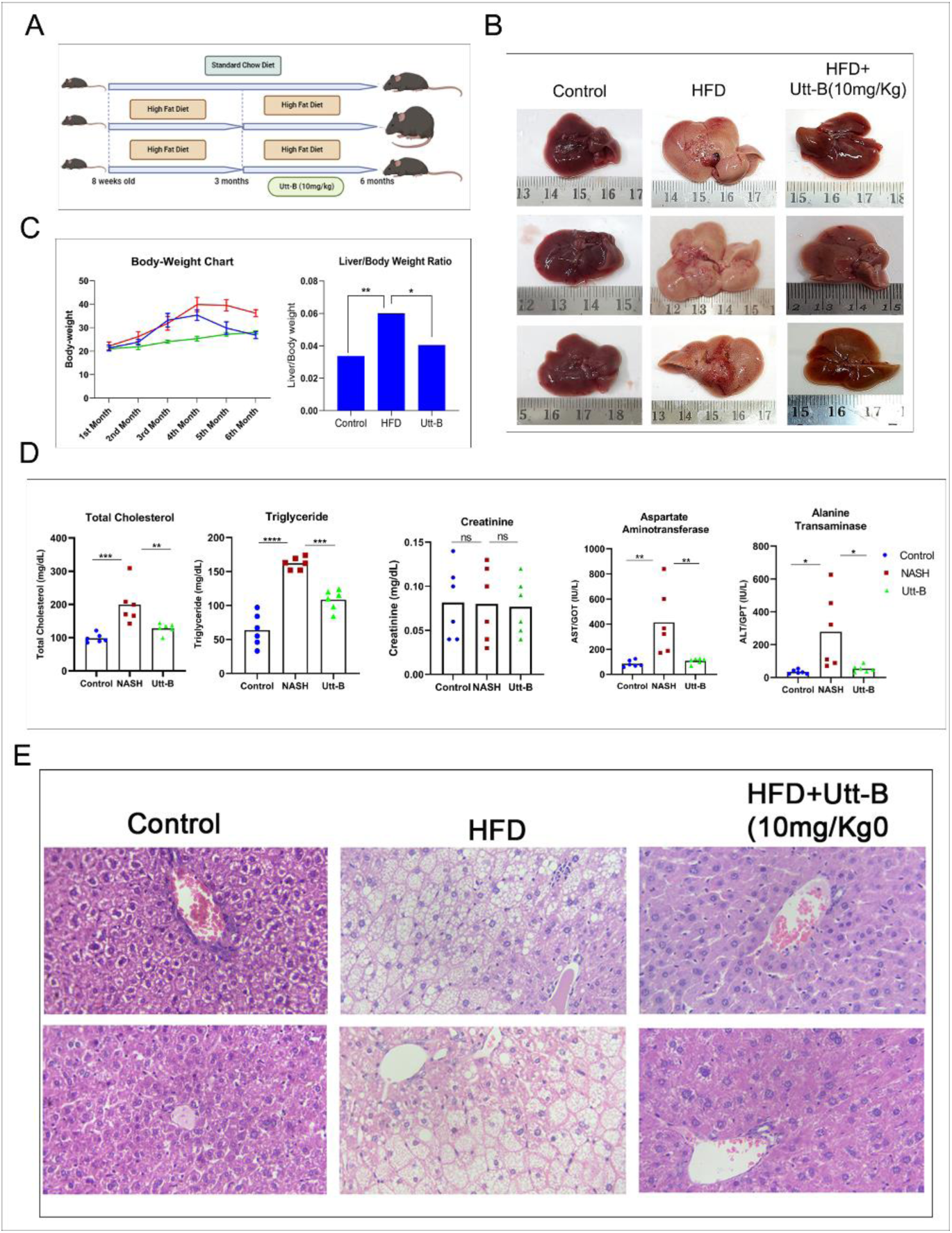
Utt-B exhibits therapeutic potential against HFD-induced NASH, *in vivo.* (A) Schematic representation of the treatment regimen followed; (B) Representative images of livers resected from Control, HFD and Utt-B-treated groups; (C) Graphical representation of Liver-Body weight ratio (D) Serum biochemical analysis of lipid profile, creatinine and liver profile indicating pharmacological safety and therapeutic efficacy of Utt-B; One-way ANOVA was used for statistical analysis; *p<0.1, **p<0.01, ***p<0.001, ****p<0.001 (E) Histopathology results depicting the development of characteristic features of NASH in HFD animals and alleviation of NASH in the Utt-B treated animals.

Serum biochemical analysis reports showed an elevated level of total cholesterol, triglycerides, ALT, AST and ALP in the HFD group, while Utt-B treated mice displayed a decrease in the serological parameters, demonstrating a trend comparable to mice fed with the standard chow diet. Renal profile analysis indicated minimal variations among the three groups, underlining the pharmacological safety of Utt-B (Fig 1D).

To assess the development of NASH in the experimental animals, histopathology analysis, which is considered as the gold technique for NASH diagnosis, was conducted (Fig 1E, S1B). Animals in the HFD group exhibited distinctive pathologic features such as, macro and micro-vesicular steatosis, ballooning of hepatocytes and lobular necroinflammation, which are conditions specifically associated with NASH, confirming the development of human-like NASH in the HFD mice (Fig S1C). Conversely, mice treated with Utt-B displayed a histologically normal liver architecture with mild steatosis, demonstrating the therapeutic efficacy of Utt-B in halting the pathological changes associated with NASH (Fig 1E, S1D).

### Utt-B inhibits inflammatory markers associated with NASH and induces autophagy, abrogating hepatic stellate cell activation, which leads to fibrosis

NASH is a condition characterized by hepatic lipid accumulation, liver cell injury and inflammation, which leads to hepatic fibrosis. Hepatocyte damage triggers inflammation and initiates a cascade of events that culminate in liver fibrosis through the activation of hepatic stellate cells (HSCs) and subsequent secretion and deposition of extracellular matrix (ECM). Hence, we investigated the pathological features of NASH, regulated by Utt-B, in the livers of the experimental animals (Fig1C). The NASH-induced mice displayed excessive lipid droplet accumulation within hepatocytes, accompanied by a significant up-regulation in the mRNA expression of the inflammatory markers, *MCP-1, CRP-1, IFN-γ* and *TNF-α,* which are associated with NASH. Remarkably, treatment with Utt-B exerted a notable reduction in lipid droplet deposition and hepatic expression levels of the inflammatory markers, demonstrating that Utt-B exhibits anti-steatotic and anti-inflammatory potential, which leads to the alleviation of NASH (Fig 2A, B).

**Figure 2:**
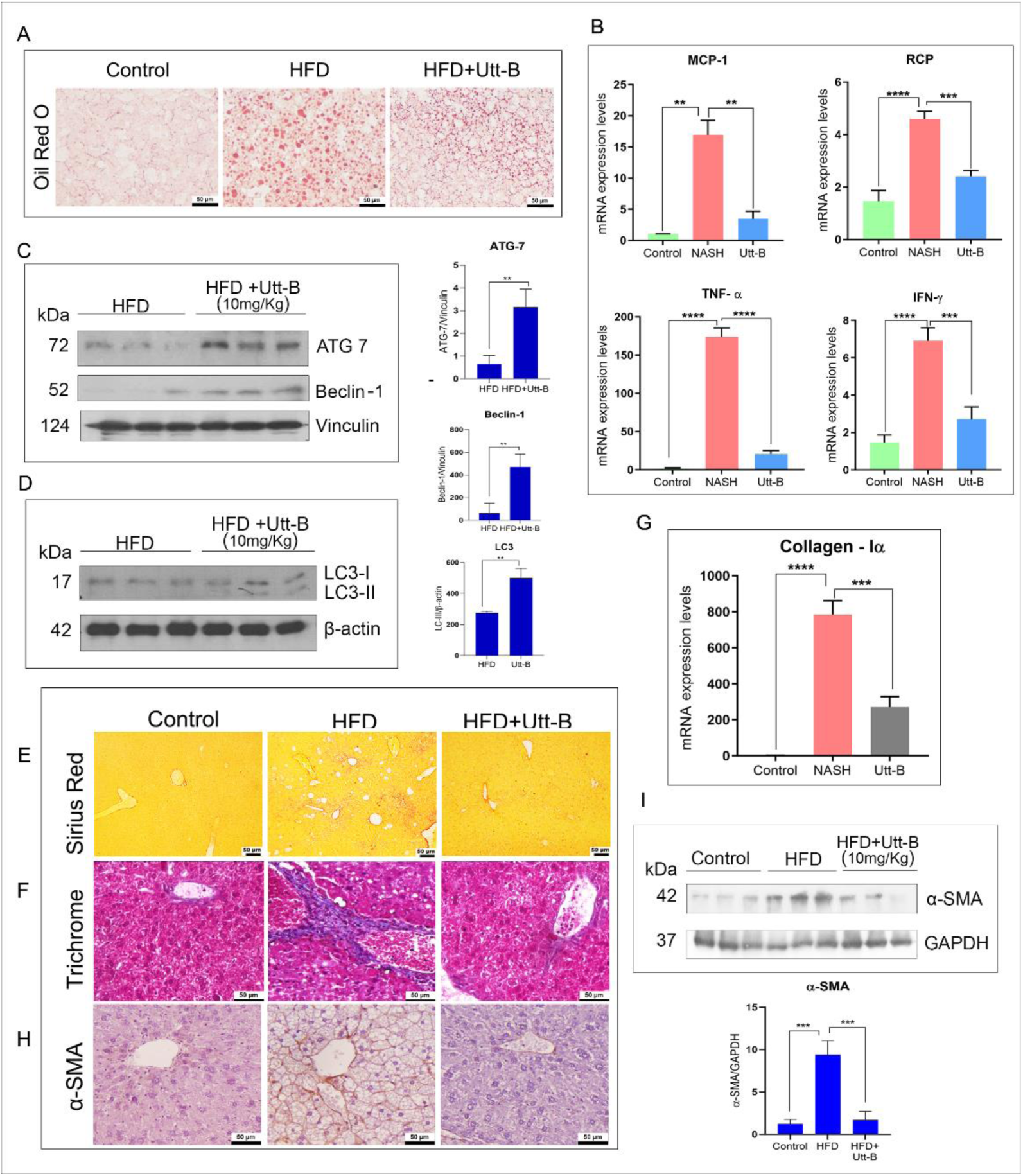
Utt-B abrogates the progression of NASH to fibrosis (A) Utt-B treatment reduces the deposition of lipid droplets within hepatocytes (B) HFD animals display an elevation of NASH-associated inflammatory markers MCP-1, TNF-α, IFN-γ and RCP, whereas Utt-B treatment results in a decrease in the expression pattern of these markers; (C) Utt-B treatment increases the expression of the autophagy markers, ATG-7, Beclin-1 and (D) LC-III; unpaired t-test was used for statistical analysis; ** p<0.05 (E) Sirius Red and (F) Trichrome staining reveals the deposition of fibrotic tissues in HFD mice and its minimal deposition in Utt-B treated mice. (G) Utt-B treatment reduces the mRNA expression of Collagen 1α; One-way Anova was used for statistical analysis; *p<0.1, **p<0.01, ***p<0.001, ****p<0.001; Utt-B reduces the expression of α-SMA, an indicator of HSC activation as assessed by (H) immunohistochemical staining and (I) Immunoblotting; One-way Anova was used for statistical analysis; *** p<0.001, **** p<0.001.

Impairment of autophagy in NASH is another major contributory factor for the progression of NASH to fibrosis. Interestingly, Utt-B treatment induced autophagy in HFD-fed animals as revealed by the increased expression of indicated autophagy markers (Fig 2C,D). Concurrently, a significant reduction in the expression of the key autophagy markers, ATG-7, Beclin-1 and LC3-II was observed in the NASH-induced animals. The liver sections of NASH-induced mice displayed regions of intense collagen deposition, while those of Utt-B-treated mice exhibited a drastic reduction in the collagen deposition as assessed by Sirius Red and Trichrome staining (Fig 2E, F). These findings were substantiated by the mRNA expression pattern of Collagen 1α, the type1 collagen, in the liver tissues of mice across all three groups (Fig 2G).

The activation of HSCs and excessive production of extracellular matrix (ECM) proteins significantly contribute to the progression of hepatic fibrosis. Immunohistochemical analysis in the liver tissues of NASH-induced mice indicated an increase in the expression of α-smooth muscle actin (α-SMA), a widely recognized marker of activated HSCs, throughout the parenchyma, compared to the mice in control and treatment groups (Fig 2H). Immunoblotting also revealed an up-regulation of α-SMA in NASH-induced mice and its down-regulation in Utt-B treated mice, illustrating the potential of Utt-B to alleviate the pathological features of NASH and attenuate the process of fibrogenesis in HFD mice (Fig 2I). These results demonstrate that the anti-lipogenic and anti-inflammatory effects of Utt-B, aid in the mitigation of the characteristic NASH features in a HFD-induced mouse model.

### Utt-B blocks the progression of NASH to HCC

We extend our investigations to inquire whether Utt-B can prevent the progression of NASH to HCC. To address this conjecture, we developed the STAM model, which employs diabetes-induced NASH, which slowly progresses to HCC, as detailed in the ‘Materials and Methods’. As per previous reports [21–23], in this mouse model, NASH develops from week 6-8, fibrosis from week 10-12 and ultimately results in HCC by the 20^th^ week. We conducted the study for a period of 22 weeks, initiating Utt-B treatment from the 8th week and continuing until the 22^nd^ week (Fig 3A). Upon euthanasia, well-developed solid tumors were observed in the livers of male mice in the STZ+HFD group whereas no tumor formation was observed in the livers of male mice that received Utt-B (Fig 3B). Studies have previously indicated that tumor formation in the STAM model is exclusive to male mice, as female hormones interfere with the action of streptozotocin. However, in our study, female mice exposed to an extended period of HFD, until the 30^th^ week, displayed the formation of nodules and well-developed tumors in the STZ + HFD group, while no such nodules were noted in the livers of Utt-B group, which displayed the liver characteristics similar to that of mice in the control group (Fig 3B).

**Fig 3:**
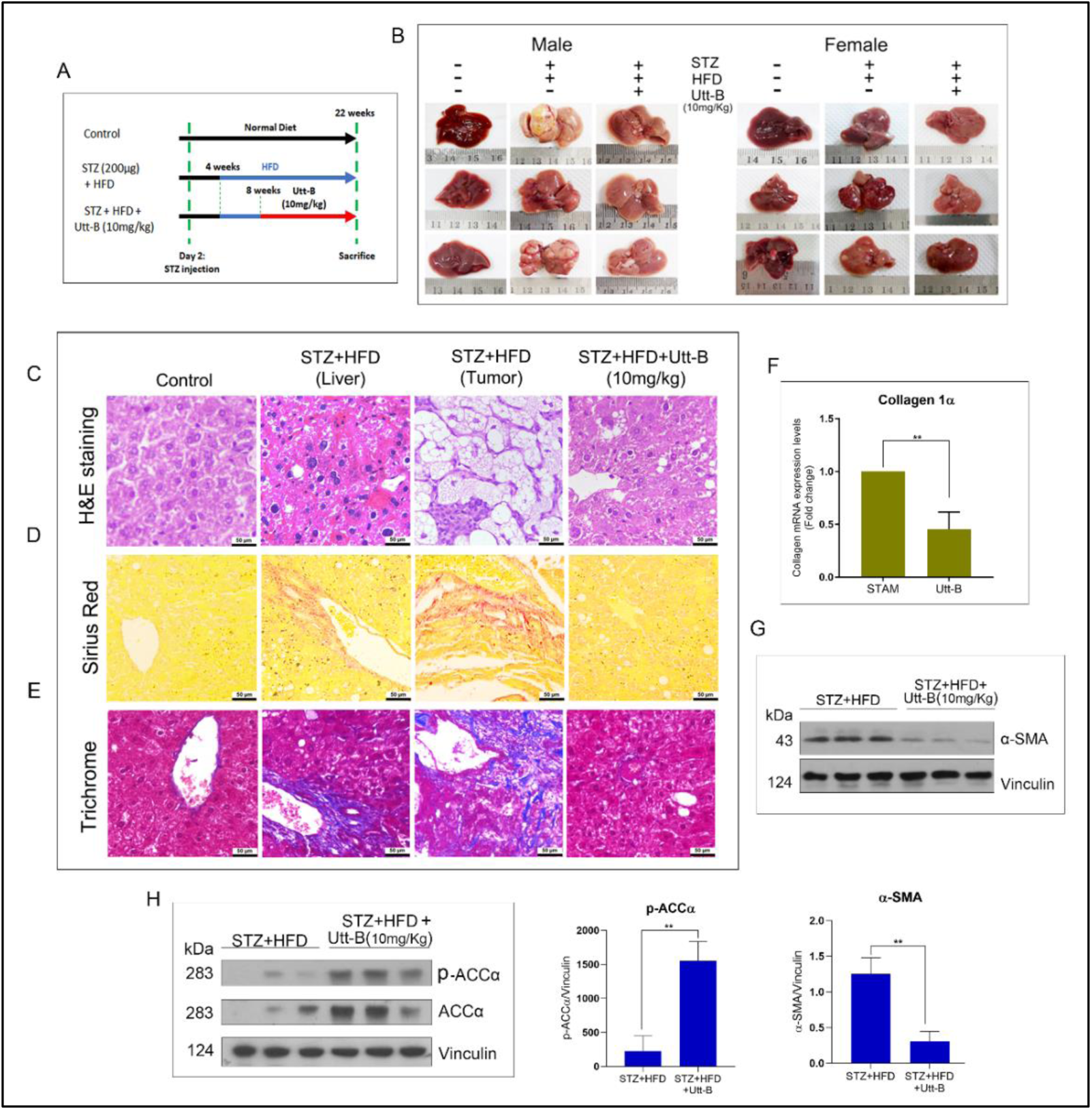
Utt-B hampers the progression of NASH to HCC: (A) Schematic representation of induction of steatohepatitis-derived HCC in BL/6 pups and treatment regimen; (B) Representative images of livers resected from male and female mice respectively, injected with sodium citrate buffer and fed normal chow diet, mice injected with streptozotocin and fed with high fat diet (STAM mice), STAM mice treated with Utt-B, indicating that Utt-B inhibits development of HCC in STAM mice model; (C) Histopathology analysis reveals formation of solid tumor in STZ+HFD group and anti-tumor efficacy of Utt-B in the treatment group; (D) Sirius Red and (E) Trichrome staining displays fibrous connective tissue in STZ+ HFD group and its alleviation on treatment with Utt-B; Utt-B reduces the mRNA expression of (F) Collagen1α; (G) induces down-regulation of αSMA; and induces the (H) phosphorylation of ACC-α; unpaired t-test was used for statistical analysis; ** p<0.05

Development of HCC in male mice within the STZ+HFD group was confirmed through histopathological analysis. Male mice in the STZ+HFD group exhibited well differentiated intranuclear HCC inclusions. In contrast, livers resected from mice in the Utt-B treatment group displayed a consistently normal histology, characterized by mild steatosis (Fig 3C). Interestingly, the female mice in the STZ+HFD group displayed an inconsistent pattern with, some developing HCC, some showing steatosis, and the others displaying a normal histology (Fig S1 E). Remarkably, all mice treated with Utt-B, regardless of gender, consistently displayed a normal histology (Fig 3C). Given the observed heterogenous pattern in female mice in the STZ+HFD group (Fig 3B, S1E), further analysis was focussed only on male mice.

Further, the development of fibrosis was compared in the liver tissues collected from male mice among the groups, by Sirius Red and Trichrome staining. The liver sections from the STZ+HFD group displayed extensive collagen accumulation, as opposed to the negligible collagen deposition observed in the Utt-B group (Fig 3D, E). This was attested by the decrease in the mRNA expression of Collagen 1α, in this group (Fig 3F). In addition, the livers of the experimental animals were also assessed for the expression of α-smooth muscle actin, an indicator of hepatic stellate cell activation. The STZ+HFD group showed an up-regulation of α-smooth muscle actin, indicating the activation of hepatic stellate cells, which could further lead to the deposition of collagen and development of fibrosis, whereas the treatment group displayed a down-regulation of the protein, pinpointing the reason behind the efficacy of Utt-B in hampering collagen deposition (Fig 3G). This observation was further confirmed by immunoblotting for acetyl carboxylase A, a rate limiting enzyme in *de novo* lipogenesis and a regulator of β-oxidation of fatty acids, which exhibits variations in its phosphorylation status during impaired lipogenesis. Interestingly, livers of Utt-B group displayed enhanced expression in the total as well as phosphorylated enzyme in comparison with the STZ+HFD group (Fig 3H). Inhibition of ACC through phosphorylation reduces the activation of HSCs, causing a reduction in α-SMA expression and collagen production and eventually suppresses the development of HCC. These results highlight the potential of Utt-B to impede the progression of fibrosis and hinder subsequent development of HCC.

### Nanostring Analysis suggests that Utt-B hinders the progression of NASH to HCC by targeting fibrosis

To elucidate the precise mechanisms by which Utt-B impedes the progression from NASH to HCC, we performed targeted transcriptome analysis of genes involved in fibrosis pathway using the Nanostring nCounter Fibrosis Panel against liver samples from both the groups. Comparative pathway score analysis revealed a marked down-regulation of fibrosis-associated pathways in Utt-B-treated group. Results from the nCounter analysis showed an up-regulation in PPAR signaling and autophagy coupled with a concomitant down-regulation of the TGF-β and PDGF signaling pathways in the Utt-B-treated group (Fig 4 A-D).

**Fig 4:**
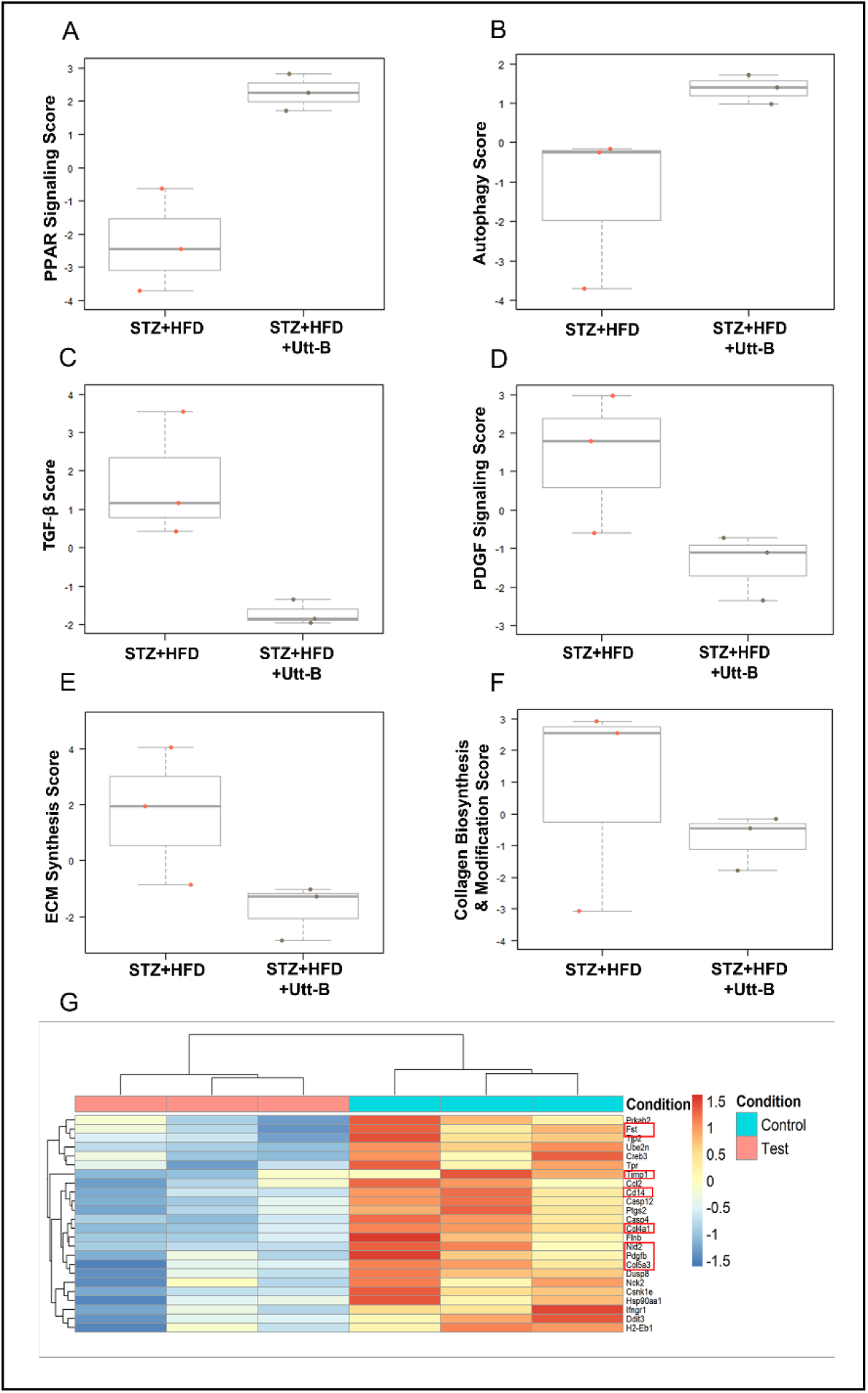
Utt-B abrogates the progression of NASH to HCC by regulating fibrogenesis: Utt-B treatment results in the up-regulation of (A) PPAR signalling and (B) Autophagy; down-regulation of (C) TGF-β (D) PDGF signalling and (E) ECM and (F) collagen biosynthesis pathways; Statistical analysis was performed using the software *R*, and P value less than 0.05 was considered statistically significant (G) Utt-B treatment results in the down-regulation of major genes involved in fibrogenesis *COL4A1, COL5A3, TIMP1, FST, CD14, PDGFB*, and *NID2*

PPARs are regarded as therapeutic targets for a variety of metabolic diseases, such as diabetes, dyslipidemia, and cardiovascular diseases, especially NAFLD. All PPAR isotypes are implicated in regulating glucolipid metabolism, inflammation, and hepatic stellate cell (HSC) activation, processes intricately linked to NAFLD. Notably, PPARγ over-expression has been reported to reverse hepatic nutritional fibrosis by curbing HSC proliferation, inducing cell cycle arrest, and promoting apoptosis. PPARγ activation has also been reported to inhibit HSC activation by down-regulating the TGFβ1/Smad signaling pathway in *in vivo* NASH model.

In our study, an up-regulation in PPAR-γ mRNA expression was observed in STZ-HFD-fed animals treated with Utt-B (S1 F). Interestingly, the most notable change identified in the comparison of gene expression between the groups was the up-regulation of genes associated with PPAR signalling (*PPARα, PPARγ* and *PGC1α*) and down-regulation of those involved in collagen and ECM biosynthesis, including *COL4A1, COL5A3, TIMP1, FST, CD14, PDGFB*, and *NID2* in the Utt-B-treated group (Fig 4G). Besides a decrease in these fibrosis markers, a down-regulation of fibrosis-related pathways such as TGF-β and PDGF pathways was also observed in the Utt-B-treated group (Fig 4C, D). Taken together, the up-regulation of PPAR signalling and down-regulation of TGF-β and PDGF pathways leading to subsequent reduction in the activation of HSCs and deposition of collagen in the Utt-B treatment group illustrates that Utt-B targets fibrosis in hindering the progression from NASH to HCC.

## Discussion

Our previous studies have demonstrated the exceptional therapeutic efficacy of Utt-B against HCC. The present study has assessed the efficacy of Utt-B in combating NASH, a critical liver disease, with an escalating mortality rate. We employed two different mouse models to study the hepatoprotective effect of Utt-B, one to evaluate the efficacy of Utt-B against NASH and the other to study its potential to hamper the advancement of NASH to HCC.

We noted a significant alleviation of the pathological features of NASH, such as lipid accumulation, macro and microvesicular steatosis, ballooning of hepatocytes and collagen deposition, by Utt-B in HFD mouse model. Interestingly, administration of Utt-B for 3 months to the NASH-developed mice, resulted in a normal liver histology with mild steatosis and the hepatic features were comparable to that of the animals fed a normal chow diet. We also observed a drastic decrease in deposition of collagen and fibrous connective tissue in the livers of the Utt-B-treated mice. This observation was attested by the corresponding down-regulation of the hepatic mRNA expression of Collagen1α in these mice. A significant down-regulation of α-SMA, an indicator of HSC activation and fibrogenesis, was also noted in the livers of the Utt-B-treated mice, highlighting the anti-fibrogenic effect of Utt-B.

Excessive triglyceride accumulation occurs in NAFLD due to imbalances in lipid metabolism and can lead to ER stress and insulin resistance [24]. This triggers the release of chemokines, cytokines and subsequent activation of HSCs leading to fibrosis [25]. In the present study, an extensive deposition of lipid droplets was observed in the livers of HFD-mice, whereas that of Utt-B treated mice displayed a drastic reduction in the size and number of lipid droplets. Subsequently, a significant down-regulation of inflammatory markers associated with NASH (MCP-1, IFN-γ, TNF-α and CRP-1) was also noted in the liver tissue extracts isolated from Utt-B treated mice. Therefore, the minimal activation of HSCs in the livers of the Utt-B treated mice in the HFD model, as indicated by a striking reduction in the expression of α-SMA and Collagen1α, could be attributed to the anti-lipogenic and anti-inflammatory action of Utt-B.

Our study also revealed a significant role for Utt-B in modulating autophagy, which has a critical role in regulating NASH. Impaired hepatic autophagy is a common feature of NASH, which contributes to hepatic lipid metabolism, ER stress and insulin resistance [26–29]. Several studies report that pharmacological and genetic interventions aimed at enhancing autophagy, improve steatosis and fibrosis in NAFLD and NASH models [29–35].

Our previous study has demonstrated induction of autophagy by Utt-B in liver cancer using *in vitro* and *in vivo* models [14]. In the present study, the livers of HFD-fed mice manifested a down-regulation of autophagy. Interestingly, a significant up-regulation of the autophagy markers, ATG7, Beclin-1and LC3-II was noted in the livers isolated from HFD-mice treated with Utt-B, indicating that induction of autophagy by Utt-B may have a direct correlation with the efficacy of Utt-B in inhibiting HFD-mediated development of NASH.

NASH-induced HCC is emerging as a prevalent cause of HCC on a global scale, warranting serious attention and comprehensive intervention strategies. The anti-NASH potential of Utt-B inspired our inquiry into its efficacy in impeding progression of NASH to HCC. For this, we used the STAM model, which employs administration of a combination of streptozotocin and high fat diet, which will eventually develop HCC within 22 weeks. While all male mice in the control group developed liver tumours within 22 weeks, none of the mice in the treatment group, which received Utt-B treatment from 8 weeks onwards, developed tumors attesting the efficacy of Utt-B, not only in ameliorating the features associated with NASH, but also in hampering the advancement of NASH to its more adverse stage, HCC.

We further investigated the response of Utt-B treatment to the level of enzyme, acetyl-CoA carboxylase (ACC), which is involved in the rate limiting step of *de novo* lipogenesis that catalyzes the conversion of acetyl coenzyme A to malonyl-CoA [36]. Phosphorylation of ACC inhibits the enzyme resulting in a reduction of *de novo* lipogenesis and an increase in fatty acid oxidation. ACC inhibition has been reported to block collagen production in LX-2 HSC cells and reduce the expression of α-SMA in primary rat HSCs, and that of COL1A1 and ACTA2 in primary human HSCs [37]. Previous studies have demonstrated that pharmacological intervention with ACC inhibitors reduce steatosis in preclinical models of NASH as well as in patients diagnosed with NASH [38–40]. Moreover, it is also reported that gene level alterations that result in a reduced phosphorylation of ACC contributes to the proliferation of liver cancer cells and development of HCC and that inhibition of ACC is an effective strategy in hindering the progression to HCC [41]. Interestingly, we also observed an up-regulation of phospho-ACC-α in Utt-B treated STZ-HFD-mice.

Development of fibrosis in NASH signifies the advancement of NASH and warrants a progressive development to much severe HCC. As Utt-B treatment mitigated the pathological features associated with fibrosis in both the HFD-induced NASH model and the STAM model mice, the latter being a model designed to study the transition from NASH to HCC, we presume that the antifibrotic effect of Utt-B might be serving as the connecting link to the remarkable efficacy of Utt-B in combating NASH and NASH-induced HCC. Utt-B treatment reduced the deposition of collagen and down-regulated α-SMA, the indicator of HSC activation, in both HFD and STAM animal models, reflecting the efficacy of Utt-B against NASH and NASH-induced HCC respectively. Results of the targeted transcriptome analysis reveal that Utt-B treatment resulted in down-regulation of genes involved in collagen production and ECM biosynthesis. Comparative pathway score analysis suggests an up-regulation of PPAR signalling and a down-regulation of TGF-β and PDGF signalling pathways, which play pivotal roles in orchestrating inflammation, fibrogenesis and development of HCC [42–44]. Several studies have reported that drugs targeting PPARs display a therapeutic effect against NAFLD, NASH and fibrosis. Moreover, PPAR agonists have been reported to inhibit liver fibrosis through down-regulation of fibrosis-related genes, TGF-β1, α-SMA, CoI-α1 and through inhibition of HSC activation [41, 45–47], which are consistent with the findings of our study. Collectively, our study outcomes in both the animal models, highlight the efficacy of Utt-B not only in ameliorating NASH but also in blocking the progression of NASH to HCC by exerting anti-lipogenic, anti-inflammatory and most importantly, anti-fibrogenic effects.

## Conclusion

Taken together, NASH and NASH-induced HCC have emerged as significant challenges, particularly due to the limited therapeutic options available. The prospect of a drug that can effectively address NASH and hinder its progression to HCC holds great promise, given that HCC stands as a leading cause of mortality in patients diagnosed with NASH. The findings of the current study present Utt-B as a promising candidate, showcasing its efficacy not only in treating NASH but also in impeding its advancement to HCC. However, additional research is needed to fully understand the mechanisms of action of Utt-B against NASH and NASH-related HCC.

## Patents

Title of the invention: Uttroside B and Derivatives Thereof as Therapeutics for Hepatocellular Carcinoma. The invention has been granted patent by the USA (US2019/0160088A1), Canada (3026426.), Japan (JP2019520425), S. Korea (KR1020190008323) and Europe (EP3463382).

## Data Availability

Raw data will be provided upon request to the corresponding author.

## Acknowledgement

The authors acknowledge the assistance provided by Ms. Devika Shaji, Ms. Shirly James, Ms. Aparna, Mr Yadu, Dr. Arya Aravind and Dr. Archana in successfully completing the work. The authors also thank RGCB Instrumentation and ARF Facility for all technical support and Theracues Innovations Pvt Ltd, Bangalore, India, a service provider for NanoString platforms, team for performing mRNA probing assays on NanoString platform and Bioinformatic analysis.

## Credit Author Statement

**Tennyson P Rayginia**: Methodology, Data curation, Investigation, Formal analysis, Software, Writing-Original draft preparation. **Chenicheri K. Keerthana**: Methodology, Data curation, Investigation. **Sreekumar U. Aiswarya**: Methodology, Data curation, Investigation. **Sadiq C. Shifana**: Methodology, Data curation, Investigation. **Jannet S**: Methodology, Data curation, Investigation. **Sanjay Suresh Varma**: Methodology. **Maria Joy P**: Investigation. **Mundanattu Swetha:** Methodology, Data curation. **Archana Payickattu Retnakumary**: Methodology, Data curation, Investigation; Visualization. **Lekshmi R Nath**: Methodology. **Kalishwaralal Kalimuthu**: Funding acquisition, Data curation. **Vishnu Sunil Jaikumar**: Methodology, Data curation. **Sankar Sundaram**: Formal analysis, Visualization, Validation. **Nikhil Ponnoor Anto**: Data curation, Writing - review & editing. **Noah Isakov**: Writing - review & editing. **Kuzhuvelil B Harikumar**: Resources, Project Administration. **Ravi S. Lankalapalli**: Resources. **Ruby John Anto**: Conceptualization, Formal analysis, Resources, Project administration, Supervision, Funding acquisition, Writing - Reviewing and Editing.

## Abbreviations

HCC: Hepatocellular Carcinoma

NASH: Non-Alcoholic Steatohepatitis

FDA: Food and Drug Administration

NAFLD: Non-Alcoholic Fatty Liver Disease

α–SMA: α – Smooth Muscle Actin

TGFβ: Transforming growth factor – β

PPAR: Peroxisome proliferator – activated receptor

FXR: Farnesoid X receptor

THR: Thyroid Hormone Receptor

GLP-1: Glucagon – like peptide 1

FGF: Fibroblast growth factor

STAM: Stelic Animal Model

CCl_4_: Carbon tetrachloride

HFD: High-Fat Diet

PBS: Phosphate Buffer Saline

ALT: Alanine transaminase

AST: Aspartate aminotransferase

ALP: Alkaline phosphatase

MCP-1: Monocyte Chemoattractant protein – 1

CRP-1: C-reactive protein 1

IFN-γ: Interferon-gamma

TNF-α: Tumour necrosis factor alpha

ATG-7: Autophagy-related 7

LC3-II: Microtubule-associated protein 1A/1B-light chain 3

STZ: Streptozotocin

HSC: Hepatic Stellate Cell

ACC: Acetyl – CoA carboxylase

PDGF: Platelet derived growth factor

PGC1α: Peroxisome proliferator – activated receptor – gamma coactivator- 1 alpha

ECM: Extracellular matrix

COL4A1: Collagen type IV alpha 1 chain

COL5A3: Collagen type V alpha 3 chain

TIMP1: Tissue inhibitor of metalloproyeinases

FST: Follistatin

CD14: Cluster of differentiation 14

PDGFB: Platelet derived growth factor subunit B

NID2: Nidogen 2

ER: Endoplasmic reticulum

LX-2: Lieming Xu-2

COL1A1: Collagen type 1 alpha 1 chain

ACTA2: Actin alpha 2, smooth muscle

CoI: α1-Collagen type I alpha 1

NIN: National Institute of Nutrition

ORO: Oil Red O

GAPDH: Glyceraldehyde-3-phosphate dehydrogenase

cDNA: Complementary DNA

qRT: PCR-Real time quantitative reverse transcription PCR

HRP: Horseradish peroxidase

DAB: Diaminobenzidine

DPX: Dibutylphthalate Polystyrene Xylene

PVDF: Polyvinyl difluoride

TBST: Tris buffer saline with tween 20

SDS-PAGE: SOodium dodecyl sulfate-Polyacrylamide gel electrophoresis

