## Supplementary material for "Uttroside B, a US-FDA-Designated Orphan Drug Against Hepatocellular Carcinoma (HCC), Impedes Non-alcoholic Steatohepatitis (NASH) and NASH -Induced HCC": Supplemental Information.docx

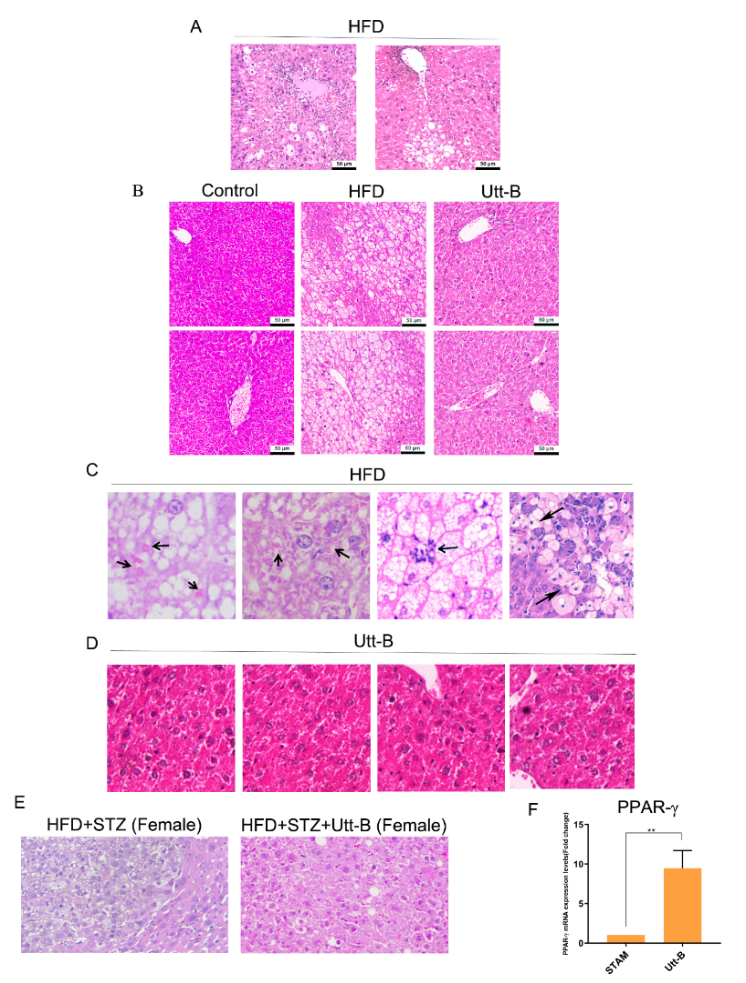

Fig S1: Supplementary Image

(A) HFD-mice exhibited steatosis and hepatocyte ballooning , the characteristic features associated with NASH, after 3 months of consumption of HFD; (B) Histopathology analysis revealed the development of NASH in HFD-mice and the alleviation of NASH in Utt-B treated mice; (C) HFD-mice displayed characteristic features of NASH such as formation of (I,ii)Mallory Denk bodies, (iii) lobular necroinflammation and (iv) hepatocellular ballooning; (D) Utt-B treated mice livers displayed a normal histology without any of the characteristic NASH features; (E) Development of HCC was confirmed in Streptozotocin-injected HFD-fed female BL/6 mice, whereas Utt-B treated mice livers displayed a normal histology with mild steatosis; (F) Utt-B treatment up-regulated the mRNA expression of PPAR-γ in steatohepatitis- induced HCC murine model
